# Tissue-Derived Insoluble Type I and Type II Collagen Inks for 3D Printing of Fibrillar Collagen Scaffolds

**DOI:** 10.64898/2026.10.29.762917

**Authors:** Hasan Rafsan Jani, Samiul Nibir, Solaiman Tarafder

## Abstract

Collagen is the primary structural component of the extracellular matrix, yet most printable collagen systems rely on soluble collagen, support baths, secondary polymers, or photochemical crosslinking. Here, we developed tissue derived insoluble type I and type II collagen inks for direct extrusion-based 3D printing without a sacrificial support bath. Type I collagen was isolated from bovine Achilles tendon and type II collagen from bovine articular cartilage. Both inks exhibited shear thinning with a predominantly elastic response that facilitated extrusion and supported shape retention after printing. The formulations were printed into grid scaffolds with infill densities of 25, 35, 40, and 50 percent and into multilayer constructs measuring approximately 20 × 20 × 20 mm. The printed constructs were treated with ammonium hydroxide to induce fibrillization and subsequently lyophilized, producing dimensionally stable and readily handleable scaffolds. Scanning electron microscopy revealed comparable porous morphologies in both scaffold types. Live and Dead staining showed abundant viable human bone marrow derived mesenchymal stem cells across the scaffold surfaces and internal regions. Cells appeared predominantly spindle shaped on type I collagen and rounded on type II collagen, suggesting distinct cell responses to the two matrices. To our knowledge, this is the first study to report 3D printable tissue derived insoluble type I and type II collagen inks within a common fabrication platform, including direct fabrication of 3D scaffolds from tissue derived type II collagen. These collagen dominant scaffolds offer promising platforms for musculoskeletal tissue engineering, regenerative grafts, and 3D in vitro models.

## 1. Introduction

Tissue engineering aims to restore lost or damaged tissue function through the integration of cells, signaling cues, and engineered scaffolds that recapitulate the structural and biological complexity of the native extracellular matrix (ECM). The ECM is not merely a passive scaffold but a hierarchically organized and dynamic network of fibrillar proteins, glycosaminoglycans, and matricellular ligands that regulate cell adhesion, proliferation, differentiation, and remodeling through coupled biochemical and biomechanical signaling (1, 2). Among its components, collagen is the most abundant structural protein in mammals and serves as a primary determinant of mechanical integrity and biological function across connective tissues (1, 3). Its triple-helical architecture, integrin-binding domains, and intrinsic ability to self-assemble into fibrillar networks make collagen uniquely suited for biomaterials design when both biological fidelity and load-bearing performance are required (1, 2, 4).

Among the 28 known collagen isoforms, types I and II are the most relevant to musculoskeletal regeneration but exhibit fundamentally distinct structural and functional roles. Type I collagen forms densely packed and highly aligned fibrils that provide tensile strength in tissues such as tendon, ligament, bone, and dermis (5, 6). In contrast, type II collagen forms a finer and highly hydrated three-dimensional (3D) network that stabilizes proteoglycan-rich matrices and supports compressive load-bearing in articular cartilage (4, 5). These structural differences are accompanied by distinct cell-instructive functions, where type II collagen promotes chondrogenic differentiation and stabilizes the chondrocyte phenotype, while type I collagen favors fibroblastic and osteogenic lineages (7, 8).

3DP has rapidly emerged as a leading fabrication strategy for spatially defined collagen scaffolds, with multiple modalities adapted for collagen-based inks (1–3). Among these, extrusion-based bioprinting enables layer-by-layer deposition of viscous inks through pneumatic or mechanical pressure and is particularly well suited for shear-thinning hydrogels that rapidly gel upon deposition (9, 10). However, the inherently poor printability of low-concentration collagen has necessitated the development of alternative strategies. Freeform reversible embedding of suspended hydrogels (FRESH) enables extrusion of acidified collagen into a thermoreversible gelatin support bath, where pH-induced neutralization triggers fibrillogenesis and stabilizes the printed architecture. This approach permits high-resolution fabrication of soft fibrillar networks that would otherwise collapse under their own weight (11, 12). FRESH-based methods have been successfully extended to anatomically scaled cardiac and cartilage constructs, to tunable microgel-bath systems for microstructural control, and more recently to the parallel printing of collagen types I, II, and III at concentrations of 20 mg/mL using commercial collagen powders (5, 13, 14). In parallel, photocrosslinkable strategies based on methacrylated collagen (ColMA) or collagen–gelatin methacrylate (GelMA) blends have been developed to achieve structural stabilization through visible or ultraviolet (UV) light mediated crosslinking, allowing tunable mechanical properties following deposition. These approaches have been further extended to digital light processing (DLP) of methacrylated type II collagen for cartilage regions and methacrylated type I collagen for subchondral regions in gradient osteochondral constructs, as well as to vat-polymerizable recombinant collagen systems (15–17).

Despite these advances, most collagen inks reported in the literature rely on acid or pepsin-solubilized type I collagen at relatively low concentrations, typically between 3 and 20 mg/mL. To compensate for limited intrinsic printability, these formulations are frequently blended with secondary materials such as alginate, hyaluronic acid (HA), gelatin or GelMA, hydroxyapatite (HAp), or xanthan gum (15, 18–22).

Although such systems have enabled the fabrication of anatomically relevant constructs and supported cell viability across a range of tissue engineering applications (12, 19, 23–28), they often compromise the native fibrillar architecture and high collagen density that are fundamental to its mechanical and biological function. These approaches, while effective for fabrication, introduce fundamental limitations. Solubilization processes disrupt the native fibrillar hierarchy and remove telopeptide-mediated intermolecular crosslinks, resulting in mechanically weak hydrogels with moduli that are often orders of magnitude lower than native tissues (5).

The low-concentration regimes imposed by solubility and viscosity constraints further limit mechanical performance and reduce the density of biologically active ligands per unit volume (9, 10, 22, 28, 29). In addition, reliance on support baths, polymer blending, or chemical crosslinking compromises the intrinsic biological identity of collagen and introduces non-native components into the system (1, 15, 18, 21). These limitations are even more pronounced for type II collagen, where ink development remains significantly underexplored despite its central role in cartilage function, with most studies restricted to cast or freeze-dried scaffold formats rather than directly printable systems (4, 7, 8, 30).

A critical and largely unresolved challenge in 3D printing of collagen is the near-universal dependence on commercially purified, acid-solubilized collagen as the starting material for ink formulation. Native connective tissues are composed of densely packed, highly organized, and largely insoluble collagen fibrils, and recapitulating this structural state requires inks that preserve rather than disrupt this architecture during processing. To date, even studies that have incorporated multiple collagen types within a single printing workflow have relied on acid-dissolved or enzyme-digested commercial powders and support-bath-based printing strategies, leaving open the question of whether tissue-derived, high-concentration, insoluble collagen can be directly processed into an extrudable ink without the need for support matrices, chemical modification, or photoinitiated crosslinking. In this study, we address this gap by developing a unified ink platform based on tissue-extracted, high-concentration insoluble type I and type II collagens derived from bovine Achilles tendon and bovine articular cartilage, respectively. Both collagens are purified through sequential chemical treatments that remove cellular components, non-collagenous proteins, proteoglycans, and lipids while preserving the native fibrillar architecture. Unlike conventional approaches, these inks are formulated at substantially higher collagen concentrations of approximately 50 mg/mL. Notably, these high-concentration, insoluble collagen inks maintain sufficient structural integrity to support large, free-standing scaffold architectures immediately after deposition, without the need for support baths or secondary stabilization strategies. This enables collagen to function as the primary structural phase rather than as a diluted or blended component. The novelty of this work lies in establishing a support-bath-free and photoinitiator-free extrusion printing framework that preserves the native fibrillar organization of collagen while enabling high-concentration processability and intrinsic shape retention.

## 2. Materials and Methods

### 2.1. Materials

Sodium chloride (S9265, Sigma-Aldrich, St. Louis, MO, USA), Tris Base (11814273001, Sigma-Aldrich, St. Louis, MO, USA), acetic acid (A6283, Sigma-Aldrich, St. Louis, MO, USA), hydrochloric acid (258148, Sigma-Aldrich, St. Louis, MO, USA), sodium hydroxide (S5881, Sigma-Aldrich, St. Louis, MO, USA), ammonium chloride (213330, Sigma-Aldrich, St. Louis, MO, USA), guanidine hydrochloride (50950, Sigma-Aldrich, St. Louis, MO, USA), ethylenediaminetetraacetic acid (798681, Sigma-Aldrich, St. Louis, MO, USA), sodium sulfate (239313, Sigma-Aldrich, St. Louis, MO, USA), ethanol, and isopropanol were used for collagen purification. Lactic acid (252476, Sigma-Aldrich, St. Louis, MO, USA) and ammonium hydroxide (221228, Sigma-Aldrich, St. Louis, MO, USA) were used for ink preparation and fibrillogenesis, respectively. Dulbecco’s Modified Eagle Medium (11885084, Gibco, Thermo Fisher Scientific, Waltham, MA, USA), fetal bovine serum (F0926, Gibco, Thermo Fisher Sci-entific, Waltham, MA, USA), penicillin–streptomycin (P4333, Gibco, Thermo Fisher Scientific, Waltham, MA, USA), and Dulbecco’s phosphate-buffered saline (14190144, Gibco, Thermo Fisher Scientific, Waltham, MA, USA) were used for cell culture. Human bone marrow-derived mesenchymal stem cells (hBMSCs) were purchased from RoosterBio (Frederick, MD, USA). The LIVE/DEAD Viability/Cytotoxicity Kit (L3224, Invitrogen, Thermo Fisher Scientific, Waltham, MA, USA) was used for cell viability assessment. All aqueous solutions were prepared using deionized (DI) water.

### 2.2. Insoluble collagen purification

#### 2.2.1. Insoluble type I collagen purification

Insoluble type I collagen was purified from bovine tendon using a modified acid and salt washing protocol based on established collagen isolation methods (31–38). Fresh bovine Achilles tendon was carefully trimmed to remove visible fat, muscle, fascia, and other connective tissues, followed by thorough rinsing with cold deionized (DI) water. The cleaned tendon was then minced into small pieces and stored at −20 °C until further processing. To remove non-collagenous proteins and soluble matrix components, the minced tissue was stirred in 0.1-0.5 M NaCl prepared in 50 mM Tris HCl buffer (pH 7.4) for 24 h. The samples were subsequently rinsed multiple times with DI water to eliminate residual salts and loosely bound impurities. The resulting insoluble, collagen-rich residue was then treated with either 0.5 M hydrochloric acid (HCl) in 0.5 M sodium sulfate (Na2SO4) for 24 to 48 h under gentle stirring. This step selectively removed acid-soluble collagen and remaining noncollagenous components while preserving the naturally crosslinked insoluble collagen fraction. Following acid treatment, the insoluble residue was separated by decanting the supernatant and extensively washing with DI water. Neutralization was carried out by immersing the material in 0.1-0.5 M NaOH under continuous stirring for 24 h, followed by repeated washing with DI water until neutral pH was achieved. To further remove residual lipids and hydrophobic impurities, the tissue was treated with 70–100% alcohol under continuous stirring and subsequently washed again with DI water. The purified collagen was finally lyophilized to obtain insoluble type I collagen as a fibrous white matrix.

#### 2.2.2. Insoluble type II collagen purification

Insoluble type II collagen was purified from bovine articular cartilage using a modified sequential salt, chelation, acid, and solvent treatment protocol adapted from established collagen isolation methods (38–42). Fresh bovine articular cartilage was dissected from femoral condyles and tibial plateaus, carefully avoiding subchondral bone and surrounding soft tissues. The harvested cartilage was thoroughly rinsed with cold deionized (DI) water to remove blood and debris, minced into small fragments, and stored at −20 °C until further processing. To remove residual blood components and loosely bound soluble proteins, the cartilage fragments were first suspended in 0.2 M ammonium chloride (NH₄Cl) under continuous stirring at 4 °C for 24 to 48 h, followed by thorough washing with DI water. The tissue was then treated twice with 0.5 M sodium hydroxide (NaOH) for 4 h to further remove cellular components and noncollagenous proteins. The resulting cartilage samples were subsequently treated with 0.5 M hydrochloric acid (HCl) containing 0.5 M Na₂SO₄ for 24 to 48 h under gentle stirring. This step was used to remove acid soluble collagen fractions and remaining non-collagenous components while retaining the highly crosslinked insoluble collagen fraction.

The insoluble material was recovered by decanting the supernatant and washed repeatedly with DI water. The cartilage samples were then treated with 0.1 M NaCl for 1 h at 4 °C to further remove soluble proteins and extracellular matrix components. After salt extraction, the tissue was extensively washed with DI water to remove residual salts. To further deplete proteoglycans and glycosaminoglycans, the samples were treated with 4 M guanidine hydrochloride (GuHCl) for 24 h under gentle stirring at 4 °C, followed by repeated washing with DI water. Chelation treatment with ethylenediaminetetraacetic acid (EDTA) was subsequently performed under continuous stirring at 4 °C to remove residual mineral components and tightly associated matrix constituents. To remove residual lipids and other hydrophobic residues, the tissue was treated with 70 to 100% alcohol under continuous stirring, followed by multiple washes with DI water to eliminate residual solvents and reagents. Finally, the purified collagen was lyophilized to obtain insoluble type II collagen as a fibrous white matrix.

### 2.3. Ink preparation from purified insoluble type I and type II collagen

#### 2.3.1. Type I Collagen Ink Preparation

Type I collagen ink was prepared by dispersing collagen powder in 0.07 M lactic acid to obtain an initial solution of approximately 0.7% (w/v). The dispersion was allowed to swell at 4 °C to ensure complete hydration of collagen fibers and facilitate uniform mixing. Following swelling, the mixture was homogenized using a high-speed homogenizer in a stepwise manner to achieve a uniform dispersion and minimize fiber aggregation. Initial homogenization was performed at moderate rotational speeds (approximately 6000–7000 rpm) to break down larger aggregates, followed by higher-speed homogenization (approximately 10,000 rpm) to further refine the dispersion and ensure a homogeneous mixture. The homogenized solution was subsequently filtered to remove undispersed particulates and then stirred continuously at 4 °C to enhance viscosity and maintain uniformity. The dispersion was further subjected to high-speed centrifugation to eliminate entrapped air bubbles and residual aggregates followed by continuous stirring at 4 °C to stabilize the ink and achieve a final collagen concentration of approximately 50 mg/mL. The resulting ink exhibited a homogeneous, high-viscosity formulation suitable for downstream applications such as extrusion-based printing.

#### 2.3.2. Type II Collagen Ink Preparation

Purified type II collagen was processed to obtain a homogeneous, high-viscosity ink through a combination of mechanical size reduction, acid-assisted swelling, blending, homogenization, and centrifugation. Briefly, lyophilized collagen was first mechanically reduced in size to facilitate dispersion. The processed collagen particles were then dispersed in lactic acid solution (0.3 M) at a concentration of approximately 1.4 wt% (w/v) and stirred magnetically at 4 °C to promote hydration and initial solubilization. Following initial swelling, the dispersion was subjected to mechanical blending in multiple cycles to break down larger aggregates and improve uniformity. The blended dispersion was returned to magnetic stirring at 4 °C for further equilibration. This sequence of blending and low-temperature stirring was repeated to progressively enhance collagen dispersion. The collagen suspension was subsequently subjected to sequential high-speed homogenization to further reduce particle size and eliminate remaining aggregates.

Homogenization was performed in multiple cycles at increasing rotational speeds (up to ∼15,000 rpm), with intermittent stirring at 4 °C between cycles to maintain uniformity and prevent thermal degradation. This stepwise homogenization approach ensured progressive refinement of collagen dispersion and improved homogeneity. Following homogenization, the collagen suspension was aliquoted and centrifuged at high speed (≈13,500 rpm) at 4 °C to remove undissolved aggregates followed by continuous stirring at 4 °C to allow gradual development of viscosity and structural organization. During this period, the collagen dispersion progressively transitioned into a highly viscous, gel-like state, indicating the formation of a concentrated collagen network. To ensure uniformity, the gelled solution was subjected to an additional centrifugation step to remove any remaining particulates. As the viscosity increased, magnetic stirring became impractical; therefore, the gel was manually collected and further processed to obtain a uniform, particle-free formulation with a final collagen concentration of approximately 50 mg/mL. The final product was a homogeneous, high-viscosity type II collagen ink suitable for extrusion-based printing and related applications.

### 2.4. Rheological Characterization of Type I and II Collagen Ink

The rheological properties of Type I and Type II collagen inks were evaluated prior to scaffold fabrication to assess printability, viscoelastic behavior, and structural stability. All rheological measurements were performed using a rotational rheometer (AR2000ex, TA Instruments, USA) equipped with a parallel plate geometry (40 mm diameter stainless steel plate). Ink samples were carefully loaded onto the lower plate, and the upper plate was adjusted to a gap of approximately 0.5 mm. Excess material was trimmed to ensure uniform contact and to minimize edge effects. All experiments were conducted at 25 °C using a temperature-controlled stage to capture relevant gel behavior for printing at room temperature. Steady shear viscosity measurements were performed by applying a shear rate ramp from 0.01 to 1000 s⁻¹ to evaluate shear-thinning behavior, which is critical for extrusion-based printing, where materials must flow under shear and rapidly recover viscosity post-deposition. Oscillatory frequency sweep tests were conducted to determine the viscoelastic properties of the inks. The storage modulus (G′) and loss modulus (G″) were measured at a constant strain of 4%, selected based on preliminary strain sweep tests to ensure measurements were performed within the linear viscoelastic region (LVR), where the material structure remains intact and the response is independent of applied strain. The frequency range was varied from 1 to 100 rad/s to capture the frequency-dependent viscoelastic behavior relevant to processing and physiological loading conditions. An oscillatory strain sweep was subsequently performed at a constant frequency of 1 Hz, with strain varying from 0.1% to 1000%, to identify the extent of the LVR and determine the critical strain at which structural breakdown and nonlinear behavior occur. This analysis provides insight into the deformation limits of the ink during extrusion and post-printing handling. A time sweep test was also performed to evaluate the temporal stability of the inks. The storage and loss moduli were monitored over 0-1000 s at a constant strain of 4% and frequency of 1 Hz, conditions chosen to remain within the LVR while assessing structural stability over time. This test is particularly important for understanding the ability of the ink to maintain consistency during printing and incubation. All rheological measurements were performed in triplicate to ensure reproducibility and reliability of the data.

### 2.5. Induction of Fibrillogenesis to Generate Fibrous Collagen Scaffolds

Printed insoluble collagen constructs were converted into fibrous scaffolds through base-induced fibrillogenesis using ammonium hydroxide. Immediately after printing, the scaffolds were frozen at −20 °C to preserve structural integrity and prevent dispersion or deformation upon subsequent immersion in aqueous solution. Frozen constructs were then directly transferred into 0.1 M ammonium hydroxide (NH₄OH) solution without thawing and incubated for 6-12 h at room temperature under static conditions. This treatment induced pH-driven fibrillogenesis and stabilized the collagen network into a fibrous architecture. Following ammonia treatment, the scaffolds were removed from the solution and washed thoroughly with deionized (DI) water to remove residual base and restore neutral pH. Washing was performed 5-6 times, with each wash lasting 30 min under gentle agitation. The scaffolds were then re-frozen at −20 °C and subsequently lyophilized to obtain dry fibrous collagen scaffolds with preserved macro- and microstructural features. This approach enables fibrillogenesis while maintaining the printed architecture, as the initial freezing step prevents collagen dispersion and preserves the structural fidelity of the constructs during exposure to aqueous alkaline conditions.

### 2.6. 3D Printing of Type I and Type II Collagen Scaffolds

The collagen I scaffold was designed using SolidWorks 2023 (Dassault Systemes) with dimensions of 30 mm × 30 mm × 3 mm. A solid scaffold geometry with 25%, 35%, 40%, and 50% infill was employed to balance structural integrity and porosity. The finalized design was exported as an STL file and fabricated using an extrusion-based TissueLabs bioprinter (TissueLabs, Bellinzona, Switzerland). The prepared collagen ink was loaded into a syringe fitted with a 22-gauge needle and printed at a flow rate of 30 kPa and a printing speed of 1 mm/s. To further assess the ability of the developed inks to support vertical build-up and multilayer fabrication, three-dimensional constructs measuring approximately 20 × 20 × 20 mm were also printed.

### 2.7. Scanning Electron Microscopy (SEM) Characterization of Type I and Type II Collagen Scaffolds

The microstructural morphology of Type I and Type II collagen scaffolds was characterized using scanning electron microscopy (SEM). Scaffold samples were sectioned from multiple regions to obtain representative microstructural features across the entire construct. The samples were carefully mounted onto aluminum SEM stubs using double-sided conductive carbon adhesive tabs to ensure proper adhesion and electrical conductivity. Prior to imaging, the mounted samples were sputter-coated with a thin layer of gold using a Plasma Sciences CRC-150 sputter coater to minimize surface charging and enhance image resolution. SEM imaging was performed using an ultrahigh-resolution scanning electron microscope (S-4700, Hitachi, Japan) operated at an accelerating voltage of 10 kV. Micrographs were acquired from multiple randomly selected regions across each sample to ensure representative characterization of fiber morphology, pore structure, and overall scaffold architecture.

### 2.8. Live/Dead Cell Viability Assay

Human bone marrow–derived mesenchymal stem cells (hBMSCs) were purchased from RoosterBio (Maryland, USA) and cultured according to our established protocols (**REFs**). Cells were maintained in Dulbecco’s Modified Eagle Medium (DMEM, Gibco, USA) supplemented with 10% fetal bovine serum (FBS, Gibco, USA) and 1% penicillin–streptomycin (PS, Gibco, USA). Cultures were incubated in a humidified environment at 37 °C with 5% CO₂, and the culture medium was refreshed every 2–3 days. Cells were expanded and passaged upon reaching approximately 70–80% confluence, and passage 3–4 (P3–P4) cells were used for all experiments to ensure consistency. For viability assessment, hBMSCs were seeded onto collagen scaffolds at a density of approximately 50,000 cells/cm² and cultured for 3 days under standard conditions. Prior to live/Dead cell viability assay, the culture medium was carefully aspirated, and samples were gently rinsed with Dulbecco’s phosphate-buffered saline (DPBS) to remove residual serum and non-adherent cells. Cell viability was evaluated using a Live/Dead assay kit (Thermo Fisher, USA) according to the manufacturer’s instructions. A solution was freshly prepared by diluting calcein AM (live cell indicator) and ethidium homodimer-1 (dead cell indicator) in DPBS. Specifically, 5 µL of calcein AM and 20 µL of ethidium homodimer-1 were added to 10 mL of DPBS. An appropriate volume (typically 200–300 µL, depending on scaffold size) of the staining solution was then added to each sample to ensure complete coverage. Samples were incubated with the staining solution under dark conditions at 37 °C to allow for dye uptake and fluorescence development. Following incubation, the collagen scaffolds were immediately imaged using a fluorescence microscope (**MICROSCOPE MODEL- Olympus IX71**). Live cells were identified by green fluorescence (calcein AM), while dead cells exhibited red fluorescence (ethidium homodimer-1).

## 3. Results and Discussion

### 3.1. Rheological Characterization of Insoluble Type I and II Collagen Ink

The rheological properties of insoluble Type I and Type II collagen inks were systematically evaluated to assess their suitability for extrusion-based printing (**Fig. 1**). Both inks exhibited pronounced shear-thinning behavior, with viscosity decreasing continuously as the shear rate increased from 0.1 to 1000 s⁻¹ (**Fig. 1a**). Type I collagen demonstrated consistently higher viscosity across the entire shear rate range, decreasing from approximately 1000 Pa·s at 0.1 s⁻¹ to ∼0.3 Pa·s at 1000 s⁻¹. In comparison, Type II collagen exhibited lower viscosity, decreasing from approximately 300 Pa·s at 0.1 s⁻¹ to ∼0.2 Pa·s at 1000 s⁻¹. This shear-thinning behavior is characteristic of pseudoplastic materials and is advantageous for extrusion-based printing, enabling flow under shear while maintaining structural integrity after deposition. Flow curve analysis revealed distinct differences in deformation behavior between the two inks (**Fig. 1b**). Type I collagen exhibited a rapid increase in shear stress at low shear rates followed by a plateau around ∼320–350 Pa over a broad shear rate range (∼200–1000 s⁻¹), indicative of yield-like behavior. In contrast, Type II collagen showed a more gradual increase in shear stress with increasing shear rate, reaching approximately ∼280 Pa at 1000 s⁻¹, suggesting a less structured network with lower resistance to flow.

**Figure 1.**
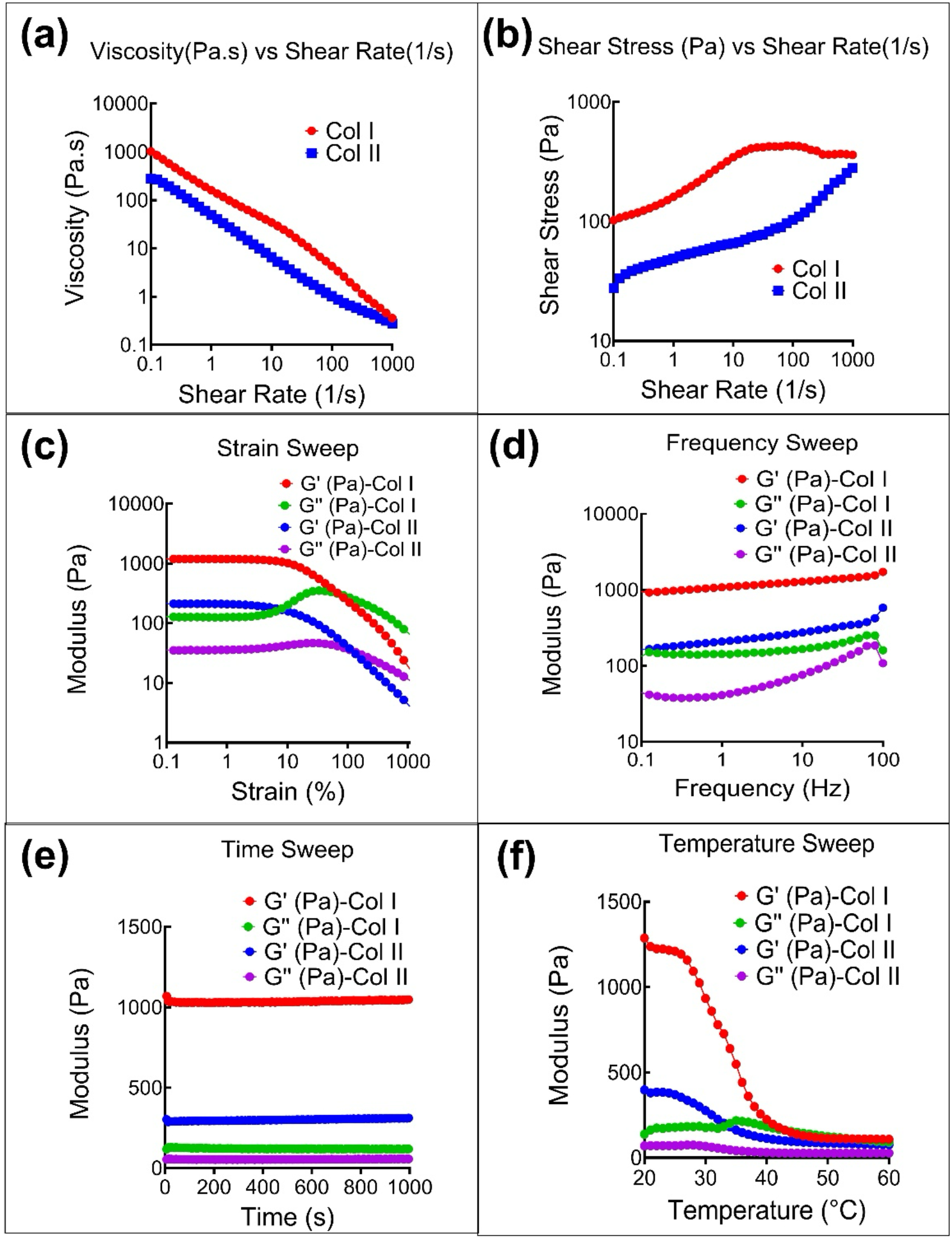
Rheological characterization of Type I and Type II collagen bioinks: Comprehensive rheological analysis of insoluble Type I and Type II collagen (Col I: red, Col II: blue) bioinks. (a) Steady shear viscosity as a function of shear rate (0.01–1000 s⁻¹) at 25 °C, demonstrating pronounced shear-thinning behavior for both bioinks, with Col I exhibiting higher viscosity across the entire shear rate range. (b) Shear stress versus shear rate profiles showing flow behavior, where Col I displays higher stress response compared to Col II, indicating greater resistance to deformation. (c) Oscillatory strain sweep showing storage modulus (G′) and loss modulus (G″) as a function of strain (0.1–1000%) at 1 Hz, identifying the linear viscoelastic region (LVR) and the onset of structural breakdown, with Col I maintaining higher moduli than Col II. (d) Frequency sweep (0.1–100 Hz) at 4% strain within the LVR, showing G′ > G″ for both bioinks, indicative of predominantly elastic, gel-like behavior, with Col I exhibiting consistently higher modulus values. (e) Time sweep (0–1000 s) at 4% strain and 1 Hz, demonstrating stable viscoelastic properties over time for both bioinks. (f) Temperature sweep (20–60 °C) showing the variation of G′ and G″ with temperature, where both bioinks exhibit a decrease in modulus with increasing temperature, indicating thermal softening, with Col I maintaining higher stiffness compared to Col II across the temperature range.

Oscillatory strain sweep measurements provided insight into the viscoelastic properties and deformation limits of the inks (**Fig. 1c**). Within the linear viscoelastic region (LVR), Type I collagen exhibited storage modulus (G′) values of approximately 1000 Pa and loss modulus (G″) values around 300 Pa, indicating dominant elastic behavior (G′ > G″). Type II collagen showed lower moduli, with G′ approximately 200 Pa and G″ approximately 50 Pa in the LVR. As strain increased, both inks exhibited a decrease in G′, indicating structural breakdown. The crossover point (G′ = G″), corresponding to the transition from gel-like to liquid-like behavior, occurred at approximately ∼200% strain for Type I collagen and ∼300% strain for Type II collagen, indicating a broader deformation tolerance for Type II collagen. Frequency sweep analysis (0.1–100 Hz at 4% strain) further confirmed the gel-like behavior of both inks (**Fig. 1d**), with G′ consistently exceeding G″ across the entire frequency range. Type I collagen maintained relatively stable moduli, with G′ ranging from approximately 1000 to 1500 Pa and G″ from 150 to 200 Pa. Type II collagen exhibited moderate frequency dependence, with G′ increasing from approximately 200 Pa at low frequencies to 500 to 600 Pa at higher frequencies, and G″ ranging from 30 to 130 Pa. The consistent dominance of elastic response (G′ > G″) indicates that both inks possess sufficient structural integrity to maintain printed shape following extrusion.

Time sweep measurements at constant strain (4%) and frequency (1 Hz) over 1000 s (**Fig. 1e**) demonstrated that both inks maintained stable storage and loss moduli over time, indicating minimal structural evolution and confirming their temporal stability under constant deformation conditions. Type I collagen maintained steady G′ values around 1050 Pa and G″ around 100 Pa, while Type II collagen exhibited stable G′ values of approximately 300 Pa and G″ around 50 Pa, with minimal variation over time. The absence of significant modulus decay or increase over time indicates that neither ink undergoes appreciable network breakdown, slippage, or time-dependent restructuring under the applied conditions. This behavior is critical for extrusion-based printing, as it ensures consistent rheological performance, sustained filament stability, and reliable shape fidelity during prolonged printing processes.

Temperature sweep analysis (20–60 °C) revealed distinct temperature-dependent behavior for both inks (**Fig. 1f**). Type I collagen exhibited a pronounced decrease in storage modulus, with G′ decreasing from approximately 1300 Pa at 20°C to around 100 Pa at 60 °C, indicating significant thermal softening. Importantly, Type I collagen maintained relatively high modulus values up to ∼30 °C, reflecting preservation of its fibrillar network and physical crosslinking interactions within this temperature range. In contrast, Type II collagen showed a more gradual decline in modulus, with G′ decreasing from approximately 400 Pa at 20 °C to around 50 Pa at 60 °C. Notably, Type II collagen also retained appreciable modulus values up to ∼30 °C, indicating sustained network integrity despite its comparatively lower stiffness. Both inks showed similar trends for G″. From a mechanistic standpoint, the retention of high modulus values up to ∼30 °C in both inks suggests that fibrillar entanglement and intermolecular interactions remain largely intact below this temperature, while progressive weakening at higher temperatures is likely associated with disruption of these physical interactions and partial network relaxation. This behavior is particularly advantageous for extrusion-based printing, as it enables processing at room temperature while maintaining sufficient stiffness and shape fidelity.

### 3.2. 3D Printing of Type I and Type II Collagen Scaffolds

Both type I and type II collagen inks were successfully extruded into grid-patterned scaffolds across four infill densities, 25%, 35%, 40%, and 50%, demonstrating the printability and structural versatility of the developed ink formulations (**Fig. 2**). The resulting scaffolds exhibited well-defined lattice architectures with regular, repeating pore geometry across all conditions. Printed filaments displayed uniform diameter and consistent inter-filament spacing throughout the scaffold body, indicative of stable flow behavior and reproducible extrusion under the applied printing parameters. Visual and structural assessment confirmed that both collagen inks maintained high shape fidelity during and immediately after printing, with no observable sagging, strand merging, or structural collapse. Following neutralization with 0.1 M ammonium hydroxide (NH4OH) solution, the scaffolds retained their macroscopic geometry while undergoing visible fibrillogenesis, evidenced by a translucent, gel-like appearance to an opaque, fibrous white matrix (**Video S1** and **S2**). After freeze-drying, both type I and type II collagen scaffolds preserved their overall lattice structure and pore architecture across all infill densities, yielding dimensionally stable, and readily handleable dry constructs. As shown in **Fig. 3**, both type I and type II collagen scaffolds could be manually lifted and handled without visible fragmentation or loss of their printed grid structure, further demonstrating the structural integrity of the processed constructs. The developed inks also supported the fabrication of thick, multilayer constructs measuring approximately 20 × 20 × 20 mm, which retained their overall 3D geometry after printing (**Fig. 4, Video S3 & S4**). The successful fabrication of these larger constructs demonstrates that the developed inks were not limited to thin planar lattices but could also sustain repeated layer deposition and vertical stacking, highlighting their potential for producing self-supporting collagen scaffolds with clinically relevant dimensions.

**Fig. 2.**
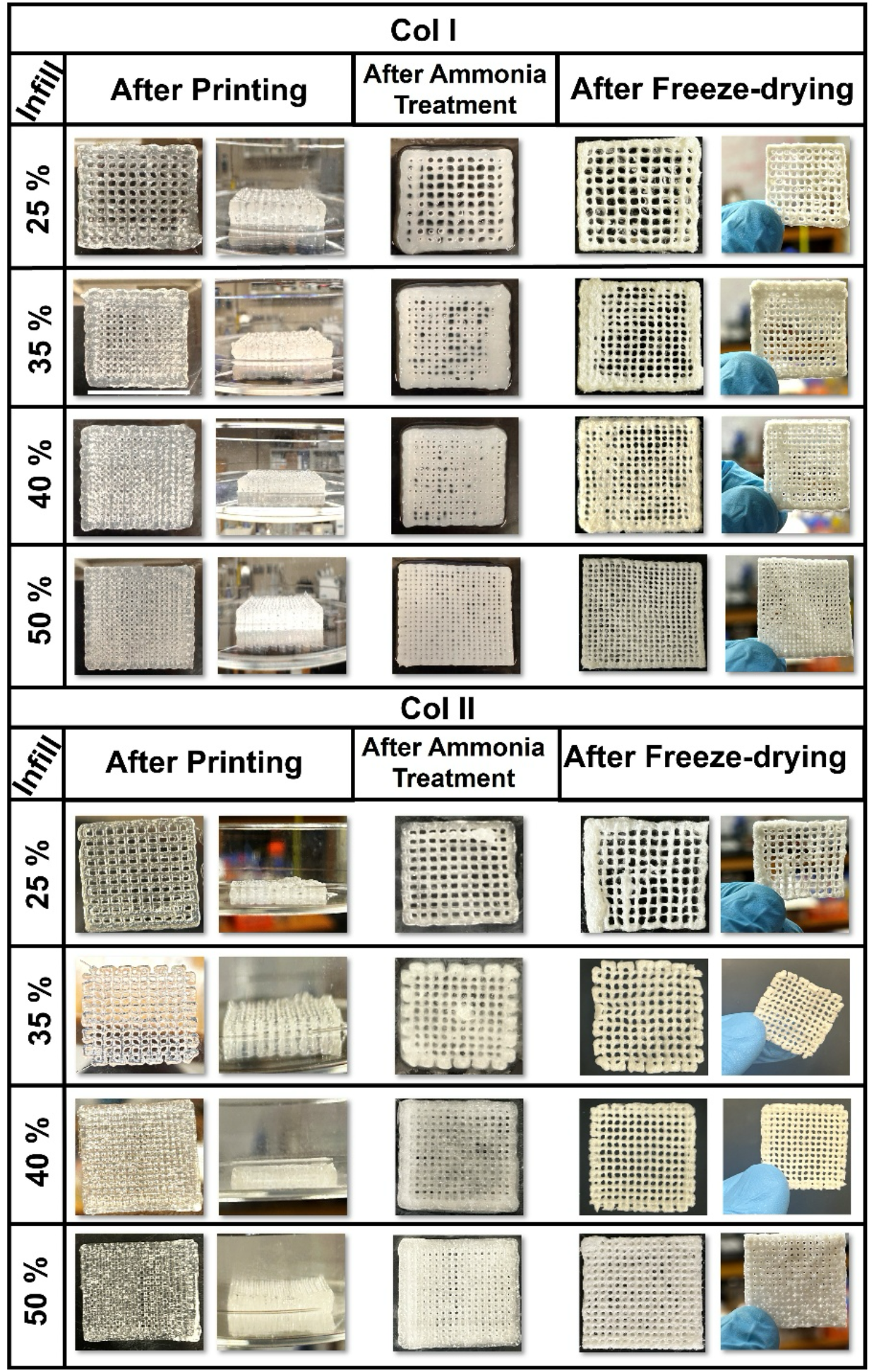
3**D Printed collagen type I and type II scaffolds at different infill densities.** Pho-tographs of successfully printed collagen scaffolds showing grid-pattern architecture at four infill densities (25%, 35%, 40%, and 50%). Type I collagen (Col I, left panel) and type II collagen (Col II, right panel) scaffolds are shown after printing, after 0.1 M ammonium hydroxide (NH4OH) treatment, and after freeze-drying for each infill condition. All scaffolds measured 30 mm × 30 mm and were printed using a 22-gauge needle at 30 kPa pressure and 1 mm/s printing speed. Images demonstrate well-defined lattice structures with regular pore g1e3ometry and uniform filament deposition across all infill densities and both collagen types.

**Figure 3.**
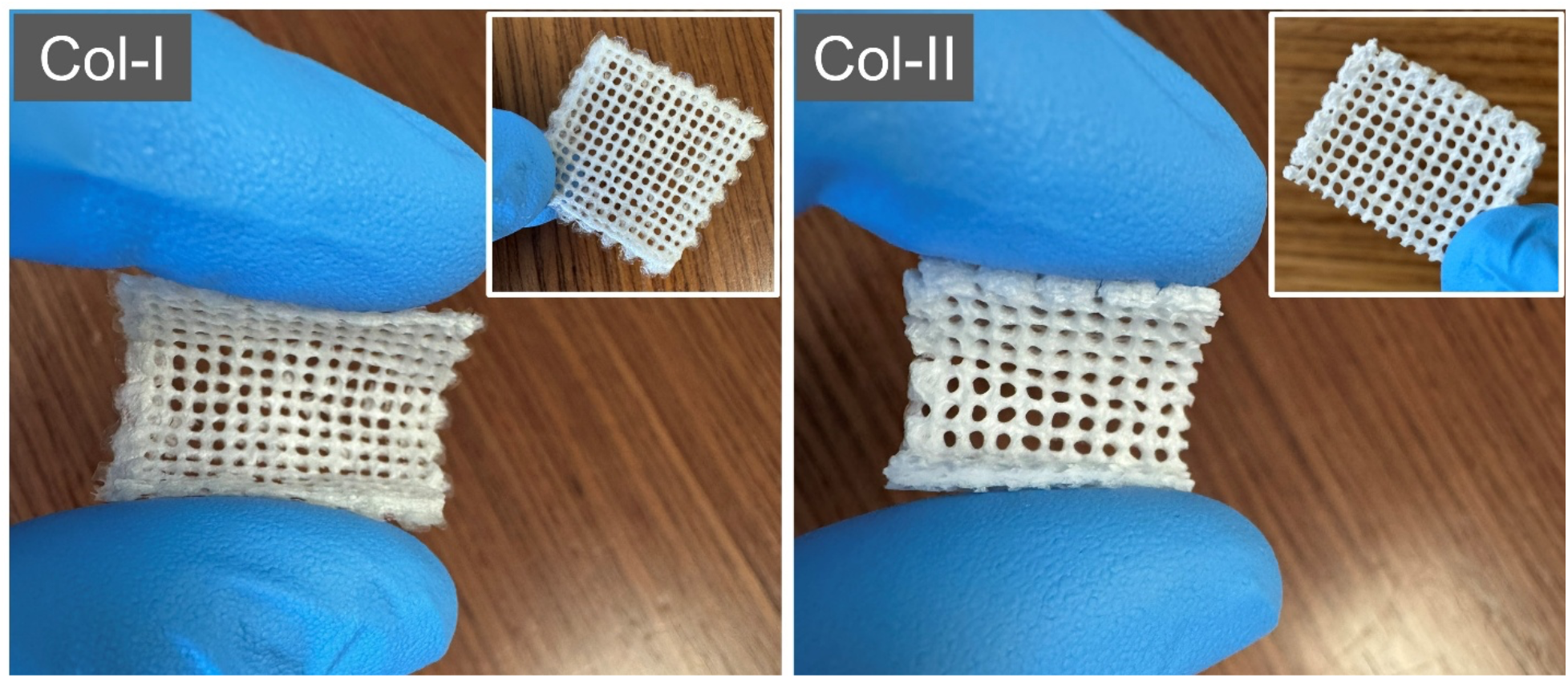
Macroscopic appearance and handleability of printed collagen scaffolds. Representative photographs of freeze-dried type I collagen (Col-I) and type II collagen (Col-II) scaffolds showing preservation of the printed lattice architecture and sufficient structural integrity for manual handling. Insets provide additional views of the corresponding grid-patterned constructs.

**Figure 4.**
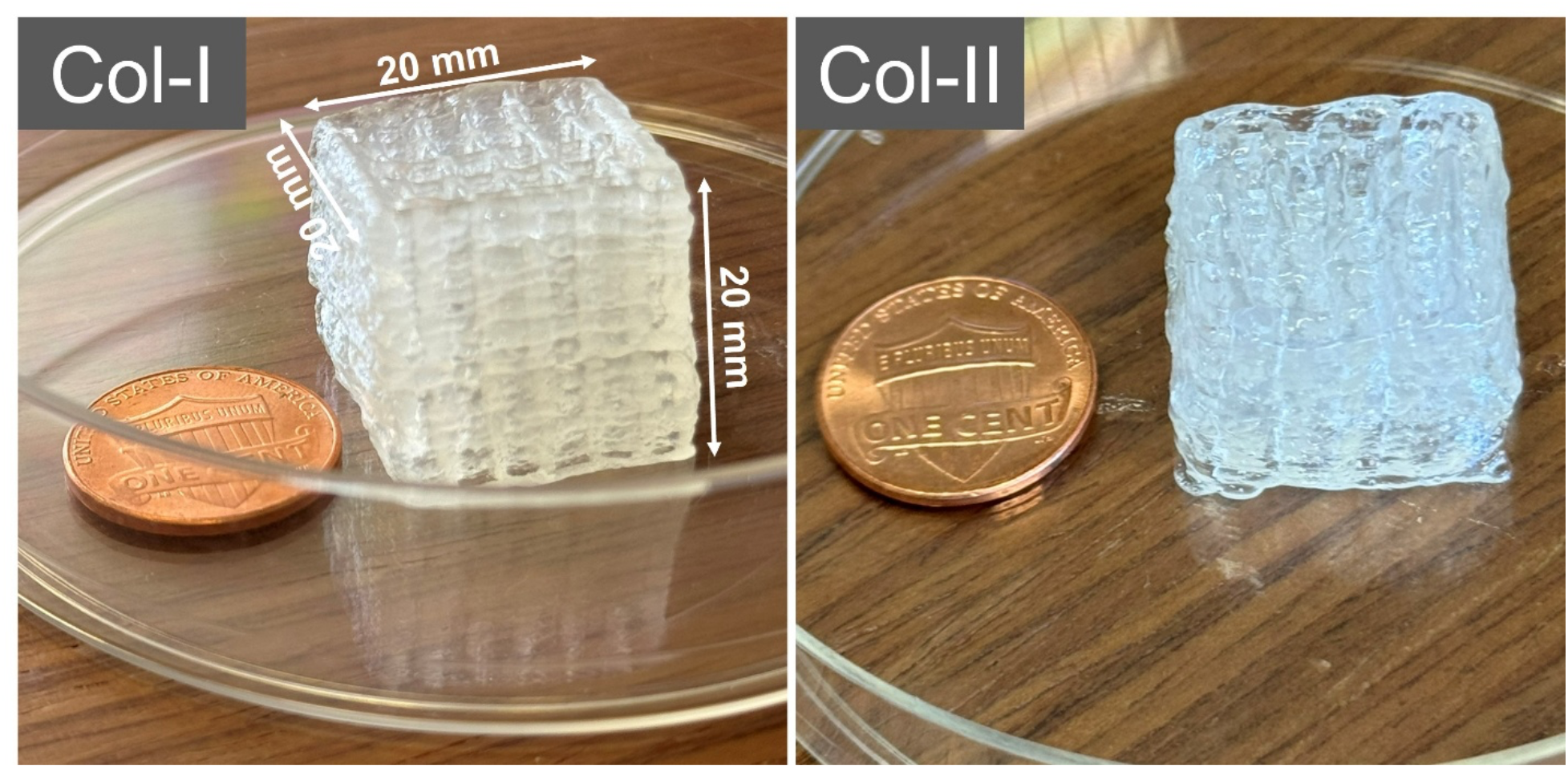
Support-bath-free printing of insoluble collagen constructs. Representative photographs of 3D printed type I collagen (Col-I) and type II collagen (Col-II) constructs as approximately 20 × 20 × 20 mm scaffolds. The coin provides a macroscopic size reference and highlights the ability of both tissue-derived collagen inks to generate self-supporting, multilayer constructs with retained overall geometry after printing.

### 3.3. Morphological Characterization of Type I and Type II Collagen Scaffolds by Scanning Electron Microscopy

SEM imaging showed that both type I and type II collagen scaffolds developed broadly comparable surface and cross-sectional morphologies after printing, ammonium hydroxide treatment, freezing, and lyophilization (**Fig. 5**). In top-surface views, both scaffold types retained clearly visible printed macropores surrounded by a continuous collagen matrix. At higher magnification, each group exhibited sheet-like regions, irregular micropores, and locally folded or corrugated matrix features. Cross-sectional images likewise revealed porous internal structures in both groups, with elongated and irregularly shaped pores distributed throughout the scaffold thickness. Both type I and type II collagen scaffolds displayed layered regions and thin internal walls consistent with structural organization induced during freezing and subsequent lyophilization. Although minor local variations in pore shape and wall thickness were observed, these differences were not consistent or sufficiently pronounced to support a clear distinction between the two collagen groups. Overall, the SEM images indicate that the two formulations produced similar porous microstructures under the applied processing conditions. The dominant morphological features appear to reflect the common printing and post-processing workflow rather than collagen-type-specific differences.

**Figure 5.**
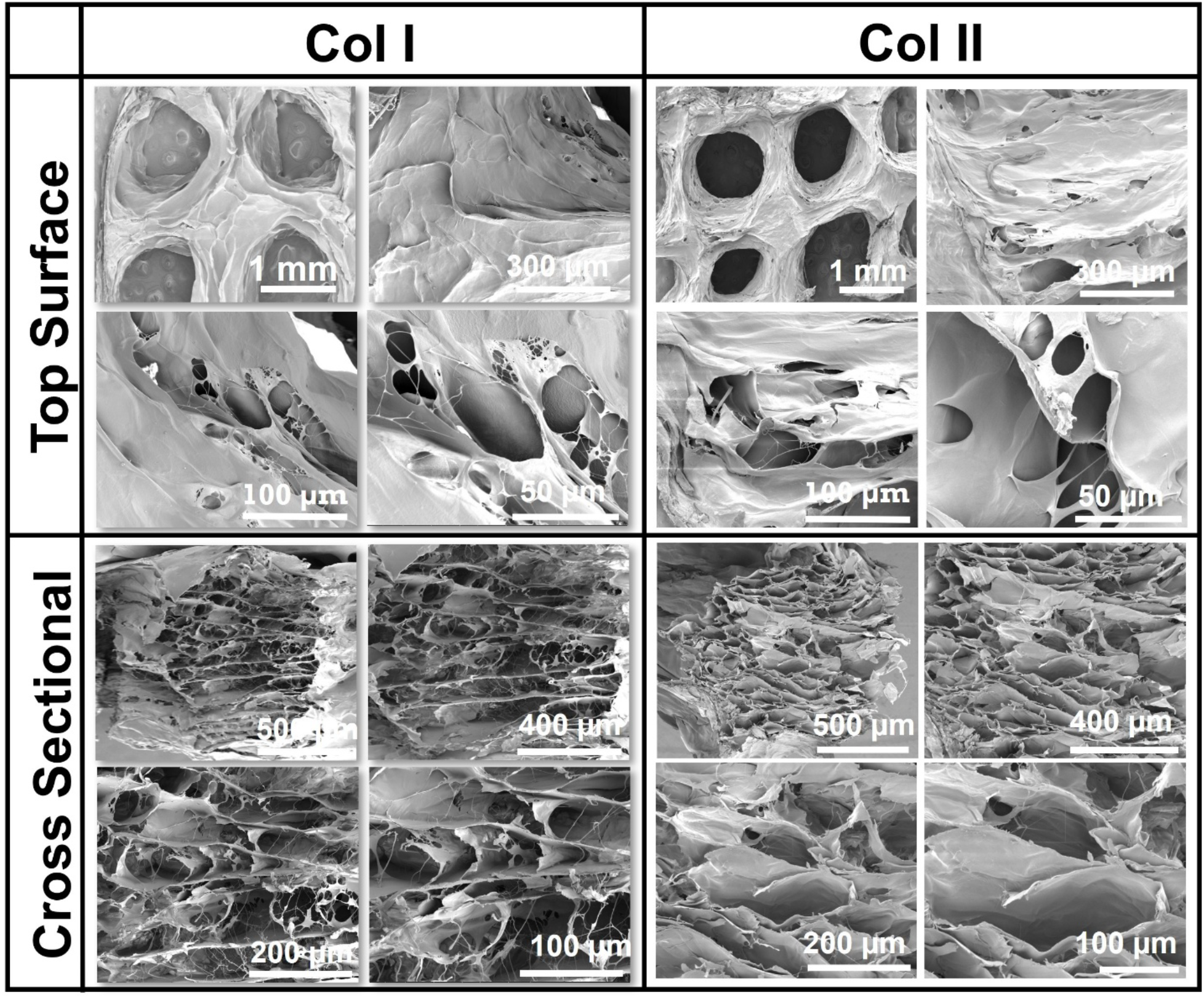
Morphological characterization of type I and type II collagen scaffolds by scanning electron microscopy. Representative SEM images of type I collagen (Col I, left columns) and type II collagen (Col II, right columns) scaffolds acquired from the top surface and cross section at multiple magnifications. Both scaffold types retained the printed macropore architecture and exhibited broadly comparable sheet-like, porous, and locally corrugated microstructural features, with no consistent morphological differences evident between the two groups.

### 3.4. Live/Dead Cell Viability Assay

The cytocompatibility of type I and type II collagen scaffolds was qualitatively evaluated by Live/Dead fluorescence staining of human bone marrow-derived mesenchymal stem cells (hBMSCs) after culture (**Fig. 6**). Both scaffold types exhibited abundant calcein AM-positive cells, evidenced by extensive green fluorescence across the printed filaments, pore regions, and deeper regions of the collagen matrix. The distribution of fluorescent cells beyond the outer filament boundaries suggests that hBMSCs were able to migrate into and populate the porous scaffold architecture rather than remaining confined to the external surface. No prominent populations of discrete ethidium homodimer-1-positive cells were evident in either scaffold group. The diffuse red signal observed throughout the images appeared to arise primarily from scaffold autofluorescence rather than clearly identifiable dead cells. No apparent qualitative difference in cell survival or distribution was observed between the type I and type II collagen scaffolds. Collectively, these observations indicate that scaffolds fabricated from the developed tissue-derived collagen inks supported hBMSC attachment, survival, and distribution within the matrix. The presence of viable cells within the scaffold regions further suggests that the internal pore network permitted initial cellular infiltration. The predominantly live cell populations also suggest that the purification, ink preparation, printing, and post-processing steps did not produce any cytotoxic response or compromise the biocompatibility of the collagen matrices. These findings highlight the potential of both scaffold types as cell-supportive platforms for biomedical applications, including tissue-engineered grafts, regenerative scaffolds, and in vitro culture models.

**Figure 6.**
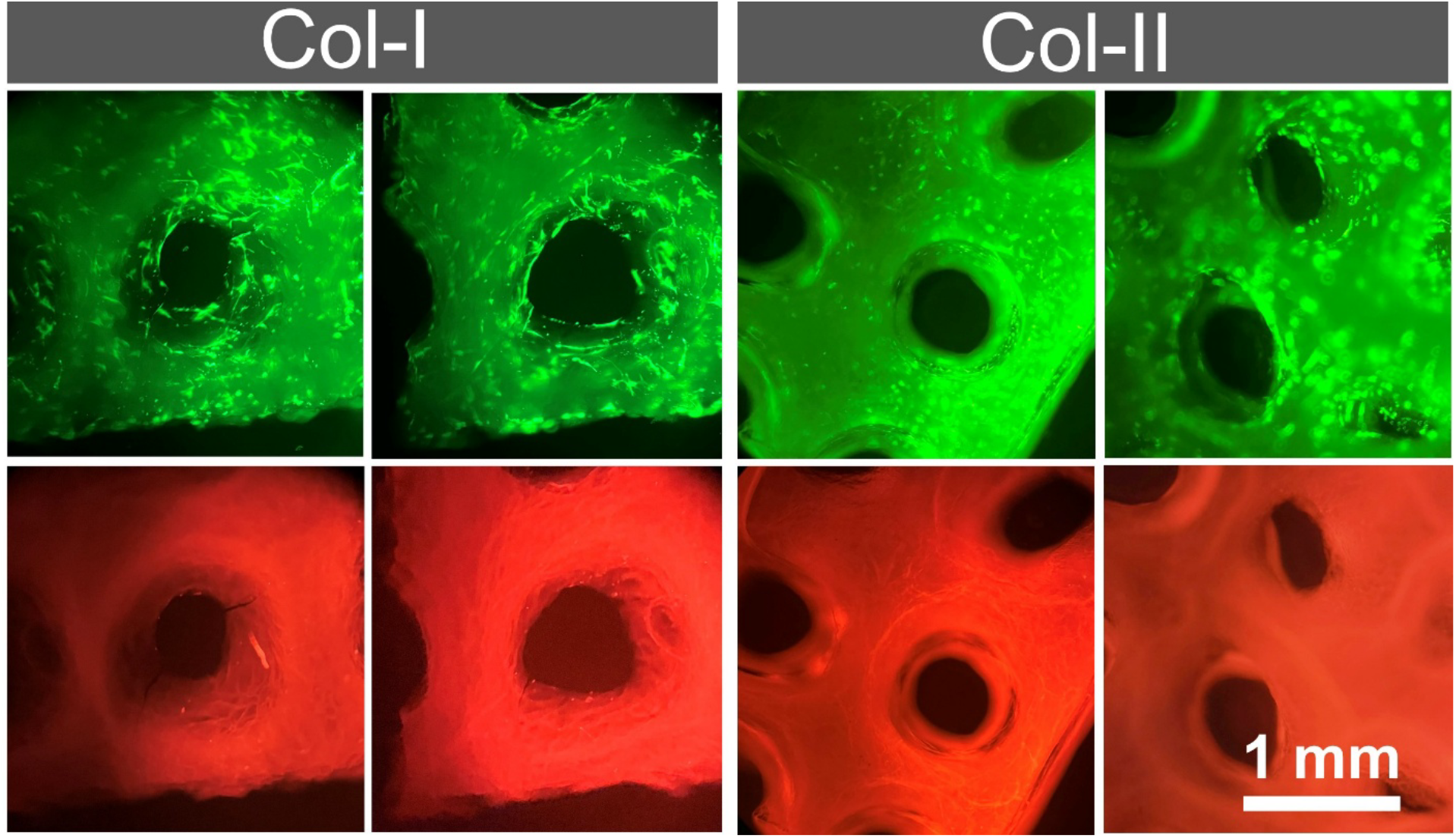
Cytocompatibility assessment of hBMSCs cultured on collagen scaffolds. Live/Dead fluorescence microscopy images showing human bone marrow-derived mesenchymal stem cells (hBMSCs) seeded on type I collagen (Col I, left column) and type II collagen (Col II, right col-umn) scaffolds following 3 days of culture. Live cells are visualized by green fluorescence (calcein AM, top row), while dead cells appear in red (ethidium homodimer-1, bottom row). Both scaffold types support predominantly live cell populations distributed throughout the scaffold filaments and inter-pore regions, with minimal dead cell signal observed beyond scaffold autofluorescence background, indicating excellent cytocompatibility of both insoluble collagen matrices.

## 4. Discussion

Insoluble collagen refers to the native, polymeric form of collagen in which the natural intermolecular crosslinks and triple-helical structure are preserved, rendering it insoluble in aqueous and dilute acidic solutions. Unlike soluble collagen (e.g., atelocollagen), which is produced through enzymatic cleavage of peptide regions that disrupts crosslinking and yields monomeric collagen molecules, insoluble collagen retains its fibrillar architecture and structural integrity. As a result, it exhibits limited solubility, higher mechanical stability, and maintains its native hierarchical organization. The novelty of this study is the development of tissue-derived insoluble type I and type II collagen ink formulations that can be extruded without a sacrificial support bath and subsequently processed into fibrillar collagen scaffolds. The two formulations, prepared at approximately 50 mg/mL, were printed as free-standing grid constructs across multiple infill densities and stabilized through base-induced fibrillogenesis while maintaining their overall macroscopic architecture. To our knowledge, this is the first study to report printable insoluble formulations of both type I and type II collagen within a common fabrication platform and their conversion into fibrillar scaffolds following printing and post-processing. Moreover, this study provides the first report of a printable tissue-derived type II collagen ink and the direct fabrication of three-dimensional scaffolds from this formulation. By avoiding secondary polymer blends and photoinitiated crosslinking, the approach maintains collagen as the dominant structural phase and establishes a distinct route for producing high-collagen-content scaffolds with intrinsic post-deposition shape retention.

The rheological response of both inks (**Fig.1**) provides a mechanistic basis for their printability. Each formulation exhibited pronounced shear-thinning behavior, enabling flow through the printing nozzle under applied pressure, while the predominance of the storage modulus over the loss modulus indicated a solid-like viscoelastic response under low-deformation conditions. This combination of shear-dependent flow and elastic character likely contributed to filament formation and resistance to immediate spreading after deposition. Type I collagen exhibited higher viscosity, shear stress, and storage modulus than type II collagen across the tested conditions, potentially reflecting differences in fibrillar organization, molecular packing, and intermolecular associations between the two formulations. Both formulations retained appreciable moduli up to approximately 30 °C, indicating sufficient network stability for processing under ambient conditions without marked thermal softening. Their moduli also remained largely unchanged over the 1000-s time sweep, demonstrating stable viscoelastic behavior under the applied low-strain conditions. Together, these findings suggest that post-deposition shape retention is supported by the inherent structural organization of the collagen inks rather than by a sacrificial support bath or externally triggered photopolymerization.

The printed constructs maintained their overall grid architecture immediately after deposition and throughout subsequent processing (**Fig. 2**). This behavior contrasts with many conventional low-concentration collagen formulations, which commonly require sacrificial support baths, polymer reinforcement, or rapid crosslinking to limit filament spreading and structural collapse (**9, 10**). Eliminating an embedding medium simplifies fabrication and avoids the additional steps associated with support-material preparation, removal, and potential entrapment within the final construct. In the present workflow, pre-freezing preserved the printed geometry during immersion in ammonium hydroxide, enabling aqueous fibrillization without substantial deformation. Ammonium hydroxide treatment neutralized the acidic collagen inks and induced fibrillar network formation (**Fig.2 and Supp Video 1 and 2**), as reflected by the transition from translucent, gel-like constructs to opaque white scaffolds while maintaining their macroscopic architecture. Thus, the combination of support-bath-free printing, freeze-assisted shape preservation, and ammonium hydroxide-induced fibrillization enabled the conversion of the printed collagen inks into structurally stable fibrillar scaffolds.

Morphological characterization by SEM showed that both type I and type II collagen scaffolds retained the printed macropore architecture and developed broadly comparable surface and cross-sectional features after ammonium hydroxide treatment, freezing, and lyophilization. Both groups exhibited sheet-like regions, irregular micropores, locally corrugated matrix features, and layered internal structures. No consistent qualitative distinction was evident between the two scaffold types. The dominant microstructural features therefore appear to reflect the shared post-processing conditions rather than collagen-type-specific assembly. In particular, the elongated and lamellar pores observed in cross section are consistent with ice-crystal templating during freezing, followed by sublimation during lyophilization.

The Live/Dead assay showed abundant viable hBMSCs distributed across the printed filaments, pore regions, and internal areas of both scaffold types. The presence of calcein AM-positive cells beyond the external filament surfaces suggests that cells were able to enter and populate the porous constructs rather than remaining restricted to the scaffold exterior. Notably, the cells on type I collagen scaffolds appeared predominantly elongated and spindle-shaped, whereas those on type II collagen scaffolds exhibited a more rounded morphology resembling the characteristic appearance of chondrocytes. This qualitative difference suggests that the two collagen matrices may provide distinct cell–material cues that influence early cell morphology. No prominent populations of discrete ethidium homodimer-1-positive cells were observed, although diffuse scaffold-associated self-fluorescence was evident in the red channel. These findings indicate that the purification, ink preparation, printing, ammonium hydroxide treatment, washing, and post-processing steps did not produce any cytotoxic response under the conditions evaluated. The ability of both scaffold types to support hBMSC attachment, survival, and initial distribution within the matrix highlights the potential of these tissue-derived collagen inks as cell-supportive platforms for biomedical applications.

The present approach complements, rather than replaces, established support-assisted collagen-printing strategies. FRESH printing has enabled high-resolution fabrication of complex and anatomically relevant collagen structures by providing temporary mechanical support during deposition and fibril formation (11, 12). More recent studies have extended embedded printing to multiple collagen types and tunable microgel support environments (5, 13, 14). In contrast, the current platform prioritizes direct extrusion of tissue-derived collagen inks without an embedding medium and uses collagen as the principal scaffold phase. Support-bath systems offer greater geometric complexity, whereas the present strategy reduces reliance on sacrificial materials and associated removal steps. Similar efforts to minimize permanent secondary phases in printed collagen constructs underscore the broader interest in simplifying collagen scaffold composition while retaining printability (43). A potential advantage of the present system is its high collagen content, which may provide a denser structural matrix than conventional low-concentration collagen hydrogels.

Overall, this study establishes a support-bath-free extrusion platform for generating fibrillar scaffolds from tissue-derived insoluble type I and type II collagen inks. The formulations exhibited rheological properties compatible with extrusion, formed free-standing grid constructs across multiple infill densities, and produced readily handleable scaffolds after post-processing. Their ability to support short-term hBMSC survival and distribution further demonstrates their promise as collagen-dominant platforms for regenerative medicine and in vitro tissue models. With additional compositional, structural, mechanical, and biological validation, this approach may provide a versatile foundation for developing tissue-specific collagen scaffolds.

## 5. Conclusions

This study establishes a direct extrusion printing strategy for tissue derived insoluble type I and type II collagen inks without the use of a sacrificial support bath, secondary polymer blend, or photoinitiated crosslinking. Both formulations exhibited rheological properties suitable for extrusion and produced free standing grid scaffolds across multiple infill densities. The inks also supported vertical layer buildup into constructs measuring approximately 20 × 20 × 20 mm. Ammonium hydroxide treatment induced collagen fibrillization and converted the printed materials into stable fibrillar scaffolds while preserving their overall geometry. After lyophilization, the scaffolds were dimensionally stable and readily handleable.

Both scaffold types showed porous internal morphologies and supported attachment, survival, and distribution of human bone marrow derived mesenchymal stem cells (hBMSCs). Cells appeared elongated on type I collagen and more rounded on type II collagen. The principal innovation is the development of a common printing platform for tissue derived insoluble type I and type II collagen inks. The work also provides the first report of a printable tissue derived type II collagen ink and the direct fabrication of 3D scaffolds from this formulation. These findings establish a promising collagen dominant platform for regenerative scaffolds, tissue engineered grafts, and 3D cell culture models.

## Supporting Information

Additional supporting information can be found online in the Supporting Information section.

## Acknowledgements

This work was supported by the National Science Foundation under the Engineering Research Initiation (ERI) Award No. 2347673 (to S. Tarafder). Any opinions, findings, and conclusions or recommendations expressed in this material are those of the authors and do not necessarily reflect the views of the National Science Foundation. This work was also supported by a Competitive Research Grant (CRG) awarded to S. Tarafder by the South Dakota Board of Regents (SDBOR).

## CRediT authorship contribution statement

Hasan Rafsan Jani: Writing – review & editing, Writing – original draft, Investigation, Methodology, Data curation, Formal analysis, Validation, Visualization. Samiul Nibir: Writing – review & editing, Investigation, Methodology, Data curation. Solaiman Tarafder: Writing – review & editing, Writing – original draft, Conceptualization, Methodology, Data curation, Formal analysis, Validation, Visualization, Resources, Supervision, Project administration.

## Data Availability Statement

Data will be made available on request.

